# Whole genome sequencing of Red Chittagong Cattle (RCC) cattle and insight into genetic variants in candidate genes for disease resistance

**DOI:** 10.1101/2023.01.09.523278

**Authors:** Ashutosh Das, Mukta Das Gupta, Mishuk Shaha, Arjuman Lima, Omar Faruk Miazi, Goutam Buddha Das

## Abstract

Detection of genome-wide genetic variation is one of the primary goals in bovine genomics. Genomes of several cattle breeds have been sequenced so far to understand the genetic variation associated with important phenotypes. Red Chittagong Cattle (RCC) is a locally adopted and disease-resistant indicine cattle breed in Bangladesh. In this study, we describe the first genome sequence of the RCC breed and in silico analyses of identified functional variants. Deep sequencing of a RCC bull genome on the NanoBall sequencing platform generated approximately 110 Gb paired-end data, resulting in 31X of genome coverage. Quality filtering retained 360,711,803 paired-end reads. Of the filtered reads, 99.8% were mapped to the bovine reference genome (ARSUCD1.2). A total of 17. 8 million Single nucleotide variants (SNVs) and 2.1 insertions and deletions (INDELs) were identified in the RCC genome. Ts/Tv ratio was computed and found to be 2.21. In total, 332 4621 variants were novel compared with dbSNP data (NCBI dbSNP bovine build 150). Functional annotation identified 54961 SNVs exonic regions, 63.75% of which were synonymous, whereas 30.42% were non-synonymous changes. The percentage of coding INDELs was 0.25% (Frameshift deletion 0.19% and Frameshift insertion 0.06%). We identified 120 variants in 26 candidates for five diseases-foot and mouth disease (FMD), Mastitis, Parasite, para-tuberculosis, and tick. Of the 120 variants, 50 were non-synonymous / frameshift (NS/FS), while 70 were synonymous/non-frameshift (SS/NFS). The identified catalog of genomic variants in RCC may establish a paradigm for cattle research in Bangladesh by filling the void and providing a database for genome-wide variation for future functional studies in RCC.

## Introduction

High throughput sequencing facilities are evolving with modern computational biological tools, and the decreasing sequencing cost enabled cost-effective variant discovery in cattle (1, 2). Furthermore, with the availability of the annotated reference bovine genome assembly and annotated genes of both taurine (*Bos taurus*) and indicine (*Bos indicus*) cattle (3, 4), comparative genomics is highly preferred for systematic genetic upgradation. Over the last decades, a substantial number of genetic variants in the form of single nucleotide polymorphism (SNP) and insertion/deletion (indel) have been identified through several bovine whole-genome sequencing studies for different cattle breeds (5). However, studies for discovering genome-wide variants are continuing since many genetic variants in diverse cattle breeds remain to be discovered.

Red Chittagong Cattle (RCC), a zebu cattle originating in the greater Chattogram district of southeastern Bangladesh, is one of the most important native cattle genotypes. RCC is found throughout the district and reared mainly under a backyard farming system with an average family of 1.6 animals. Their productive and reproductive performance (one calf per year) is better than other indigenous genotypes in Bangladesh (6, 7). Economic and genetic evaluation studies have shown that RCC farming is more profitable than other local or crossbred cattle farming under rural conditions (8-10). This genotype is quite resistant to parasites compared to other indigenous cattle in Bangladesh (6, 11-13). RCC performs well at farm and field levels (14, 15) and has a positive economic value for body weight gain (10). RCC could be considered a genotype of choice while breeding for disease resistance and beef production in Bangladesh. Most of the research works done so far on RCC have been phenotype based. Few molecular studies have been carried out to investigate RCC’s origin and genetic diversity (16-20).

However, despite past decades of research, the scientific community still needs a comprehensive database on genetic variation in the RCC because most works were descriptive. Therefore, considering the research gap and the usefulness of high throughput sequencing, we sequenced the whole genome to detect genome-wide differences between taurine and an indicine cattle breed and to reveal breed-specific genetic variants for subsequent use in breeding and conservation programs. In addition, genetic variants in candidate genes associated with disease resistance were also investigated to identify putative candidate mutation for a vital phenotype that defines the RCC genotype.

## Materials and methods

### Ethics statement

DNA was extracted from semen sample collected from Bangladesh Rural Advancement Committee (BRAC) artificial insemination enterprise (BRAC-AIE) and therefore no specific ethical approval is needed.

### DNA sampling, library construction and sequencing

Genomic DNA from a RCC bull was extracted using the Monarch Genomic DNA Purification Kit (New England Biolabs) according to the manufacturer’s guidelines. The bull selected for this study was one in the first group of RCC bulls used in the BRAC-AIE in the greater Chattogram division of Bangladesh and was the first-ranking bull in the phenotypic evaluation for the RCC bull used in the region (28,488 semen dose used, so far). In addition, this animal was one of the fathers of the first group of families of open nucleus breeding for RCC selection. The quality and purity of the extracted DNA were assessed using NanoDrop™ One Microvolume UV-Vis Spectrophotometer (Thermo Scientific). Purified genomic DNA was sent to Beijing Genomics Institute (16th Dai Fu Street, Tai Po Industrial Estate, Tai Po, N.T., Hong Kong) for library preparation (Short Insert library) and sequencing. The sequencing libraries construction was performed following the manufacturer’s recommendations for DNBSEQ Short-read library preparation.

Whole genome sequencing was performed using a DNBseq platform. First, adaptor sequences, contamination, and low-quality reads were removed from raw reads using SOAPnuke (21). Briefly, the first raw data with adapter or low-quality sequences were filtered. Then, data processing was performed to remove contamination and obtain valid data. SOAPnuke software filter parameters were: “ -n 0.001 -l 10 --adaMR 0.25”. Steps of filtering were: 1) Filter adapter: if the sequencing read matches 25.0% or more of the adapter sequence (maximum two base mismatches are allowed), remove the entire read; 2) Filter low-quality data: if the bases with a quality value of less than 10 in the sequencing read account for 50.0% or more of the entire read, delete the entire read; 3) Remove N: if the N content in the sequencing read accounts for 0.1% or more of the entire read, delete the entire read and 4) Obtain Clean reads: the output read quality value system is set to Phred+33.

### Mapping and variant calling

Raw reads were again checked for quality using fastqc (22), and low-quality raw paired reads were further filtered out using Trim Galore (https://github.com/FelixKrueger/TrimGalore). High-quality reads were mapped against the reference Bos taurus genome GCF_0002263795.1(4) using Burrows-Wheeler Aligner (BWA (23) software with the demo BWA mem settings. Mean coverage and breadth of reference genome coverage were estimated using samtools version 1.7 and bedtools v2.26.0. Variant calling was performed using SAM tools (24). Identified SNPs will further be screened using bcftools (25) and vcftools to obtain high-quality SNPs. Identified SNVs and INDELs were filtered according to the following criteria: (1) low-quality variants (PHRED <20) were removed; (2) minimum rate of two equal reads covering the least frequent allele and nucleotide positions with more than two alleles were removed; (3) variants with too low (<5) or too high (>100) read depths were removed, variants in the interval between the minimum of four reads of depth and the maximum three standard deviations above the mean were kept; (4) SNVs within 5 bp of each other and INDELs within 10 bp of each other were removed (Choi et al. 2013, adapted). The SNPs successfully passed filtering criteria were retained for variant annotation.

To identify the proportion of shared variations (SNVs and INDELs) between RCC and European taurine and indicine breeds, we compared SNVs identified in this study and SNVs reported in the dbSNP database. An in-house shell script was used to perform this analysis.

### Functional annotation of variants

Annovar (26) was used to assign a functional class to each SNV and INDEL and to describe the affected transcript and protein, if applicable. Ensembl release 94 databases were used for building the cattle database while performing the functional annotation. Each exonic SNV and indel were assigned with a diverse range of functional categories based on genomic coordinates, functional class, codon change, gene name, transcript biotype, gene coding, transcript ID, exon rank, and corresponding genotype. Annotation results were downloaded for further downstream analysis.

### Gene ontology (GO) analysis

PANTHER version 17.0 (27) was used to perform enrichment analysis of genes harboring high or medium-impact variants in this study (GO Ontology Database Released 2022-02-22). We used all *Bos taurus* genes in the PANTHER database as the reference list in our analysis. The gene list used in the analysis encompassed the gene symbols for 6810 genes, of which 5785 were uniquely mapped with the reference. Therefore, our enrichment analysis represents results for 5785 genes predicted to be affected by high or moderate-impact variants. We used the complete GO biological process as the annotation data set. Default settings for the binomial test with Bonferroni correction for multiple testing was executed to estimate the overrepresentation of functional categories. We also analyzed the KEGG category enrichment for the genes containing loss-of-function (LoF) variants (defined as stop codon, splice site, frame-shift and large deletions in protein coding genes).

### Identification and prioritization of variants in candidate genes for disease resistance

To investigate whether the candidate genes for disease resistance containing the homozygous nonsynonymous SNVs, stop gain, and frameshift deletions/insertions were associated with economic traits, we review the literature, particularly the reports that described the genomes of cattle breeds (28-30) and screened CattleQTLdb (http://www.animalgenome.org/cgi-bin/QTLdb/BT/index).

## Results and Discussion

### Sequencing and read mapping

Sequencing of RCC generated approximately 110 Gb paired-end data. After all processing of filtering approximately 360,711,803 (108 Gb) of high quality paired-end reads were retained for downstream analysis. The average read coverage across the genome was 31X. Mapping results show that 99.8% of the reads were mapped to the bovine reference genome (ARSUCD1.2). The average read coverage across the genome was in the same range as other deep re-sequencing studies (31-34). Mapped reads covered 98.4 % of the reference genome at 5x coverage, whereas 99.6% covered at least 1x coverage. This percentage of coverage was similar to Zwane et al. (35) but higher than Iqbal et al. (36) and Rosse et al. (37).

### Identification of SNVs and INDELs

In comparison to the taurine reference genome, we identified 17. 8 million SNVs and 2.1 million INDELs in the RCC genome. After filtering, 19469196 variants were retained in the filtered data set. Of which 8017539 were homozygous and 11451657 were heterozygous. Compared with dbSNP data (NCBI dbSNP bovine build 150) 3324621 variants (17.24%) were novel and 15963727 were found in the dbSNP build 150 (Fig 1). The percent of novel variants was lower compared to Rosse et al.(37) and Zwane et al. (35).

**Fig1.**
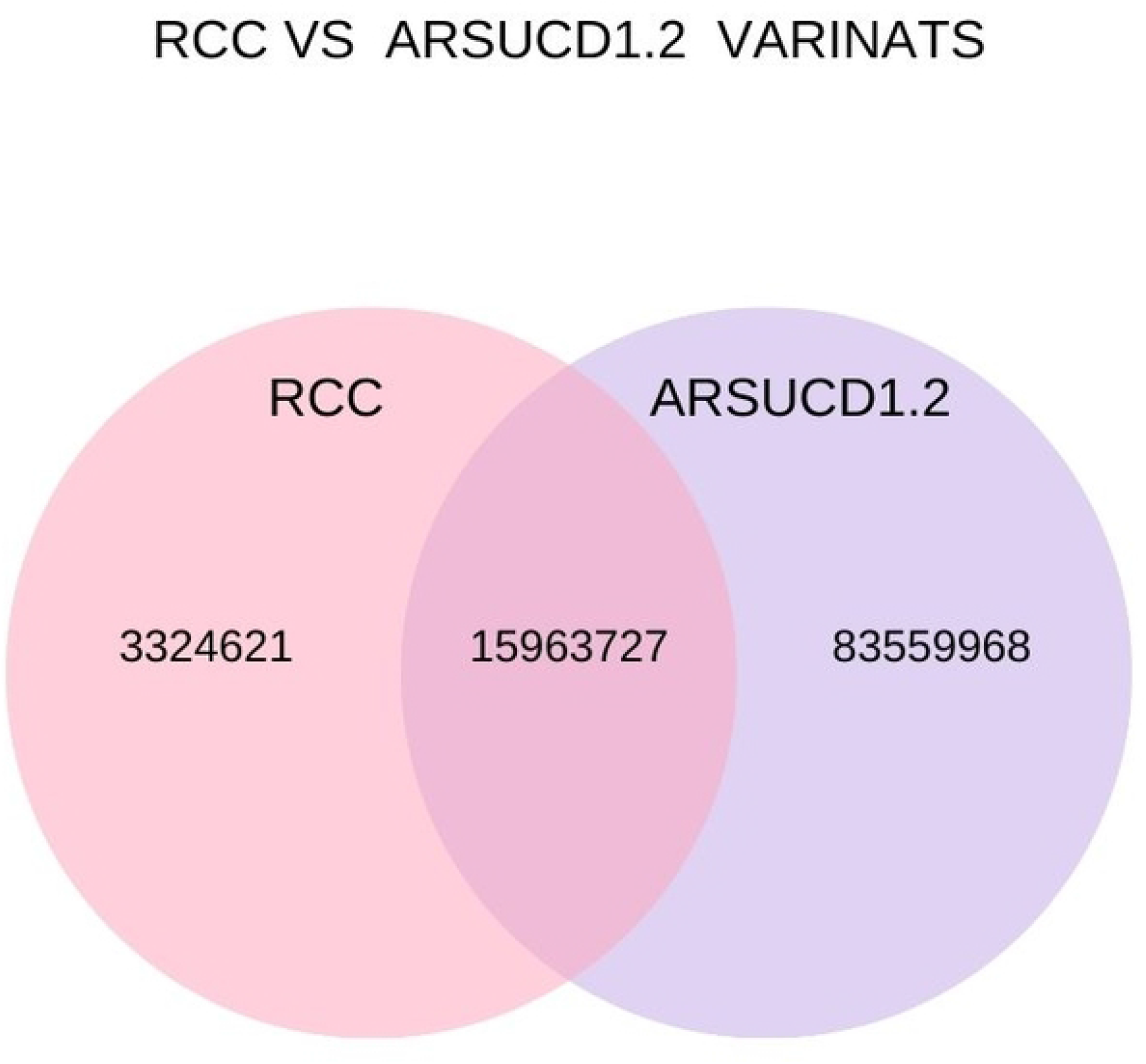
Venn diagram showing overlapping known and novel variants in Red Chittagong cattle (RCC) genome compared to taurine reference genome (ARSUCD1.2)

To evaluate the quality of the detected SNPs, the Ts/Tv ratio was computed and found to be 2.27 (Table 1). The observed Ts/Tv ratios were consistent with those observed in previous studies on different cattle breeds (31, 35, 38), indicating the quality of SNP data in the present study.

**Table 1.** Descriptive statistics of variant (SNV and INDEL) identified in Red Chittagong (RCC) cattle.

| Item | Count |
| --- | --- |
| Total variant | 19469196 |
| Number of SNVs | 17320719 |
| Number of Indels | 1967629 |
| Number of multiallelic sites | 180848 |
| Number of multiallelic SNV sites | 25191 |
| Transitions | 12048432 |
| Transversions | 5297478 |
| TS/TV | 2.27 |

### Annotation of variants

In our data, 54961 SNVs were detected in exonic regions, of which 24575 were homozygous, and the remainder was heterozygous. In total, 3824953 variants were located in intronic regions, 43589 in 3 UTRs and 7193 in 5UTR. The variants detected in upstream and downstream genes were 68337 and 58191. In addition, 15397123 (79%) variants were located in intergenic regions. Moreover, 14635 variants were annotated as ncRNA variants. The number of overlying splicing sites was 214 (Table 2).

**Table 2.**
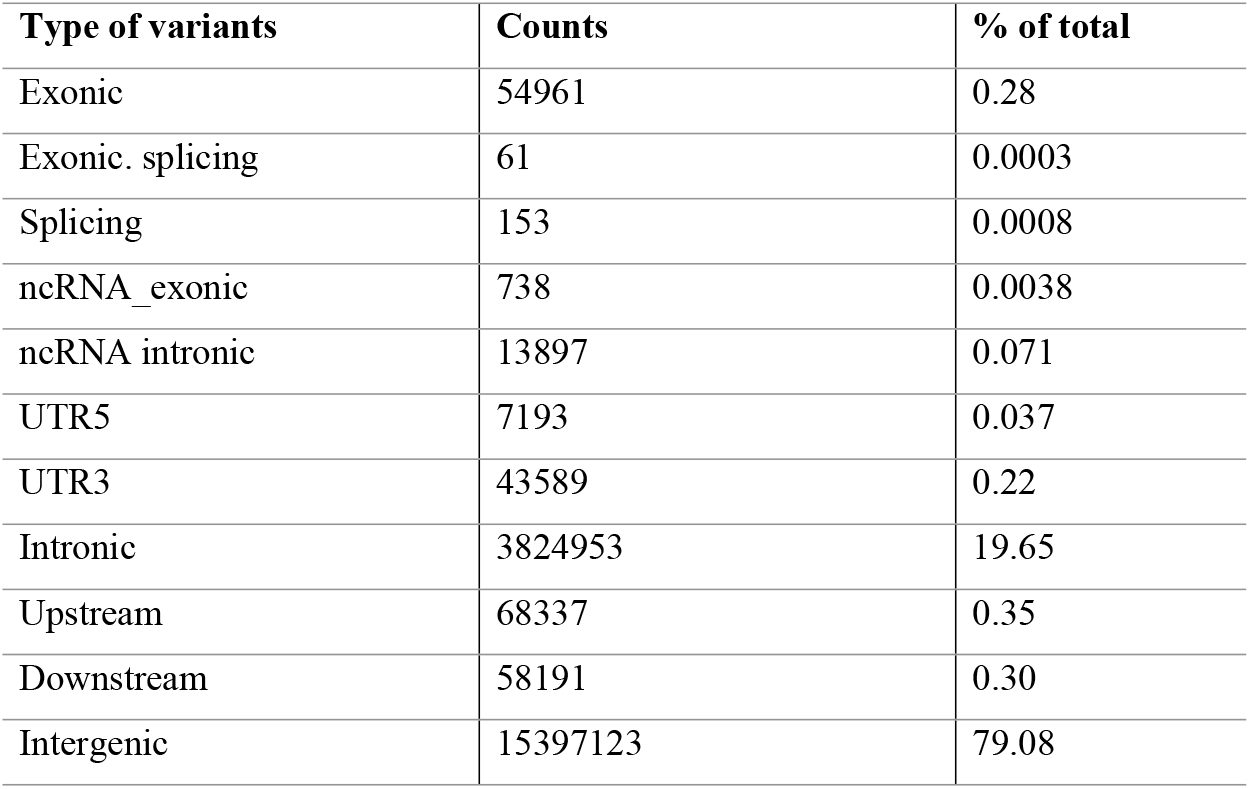
Annotation of the detected SNPs and INDELS.

The functional categories of the exonic variants are presented in Table 3. As expected, most exonic variants (63.75%) were synonymous, whereas 30.42% were non-synonymous. The percentage of deletions and insertions predicted to cause a frameshift in the protein products was 0.25%(Frameshift deletion 0.19% and Frameshift insertion 0.06%).

**Table 3.** Count of exonic variants within each functional class.

| Functional class | Number | % of total |
| --- | --- | --- |
| Synonymous SNV | 35038 | 63.75 |
| Nonsynonymous SNV | 16717 | 30.42 |
| Stop gain | 91 | 0.16 |
| Frameshift deletion | 104 | 0.19 |
| frameshift insertion | 34 | 0.06 |
| Non frameshift insertion | 158 | 0.03 |
| Non frameshift deletion | 181 | 0.29 |
| Stoploss | 16 | 0.33 |
| Unclassified | 2622 | 4.77 |

### GO Analysis of the variants

Results of GO enrichment analysis results showed 5785 known genes (out of 6810 predicted genes) harboring high or moderate impact variants belong to 20 functional categories including cellular process (29.20%), metabolic process (16.80%), biological regulation (15.10%), response to stimulus (9.30%), localization (7.40%) and signaling (6.50%) (Fig 2).

**Fig 2.**
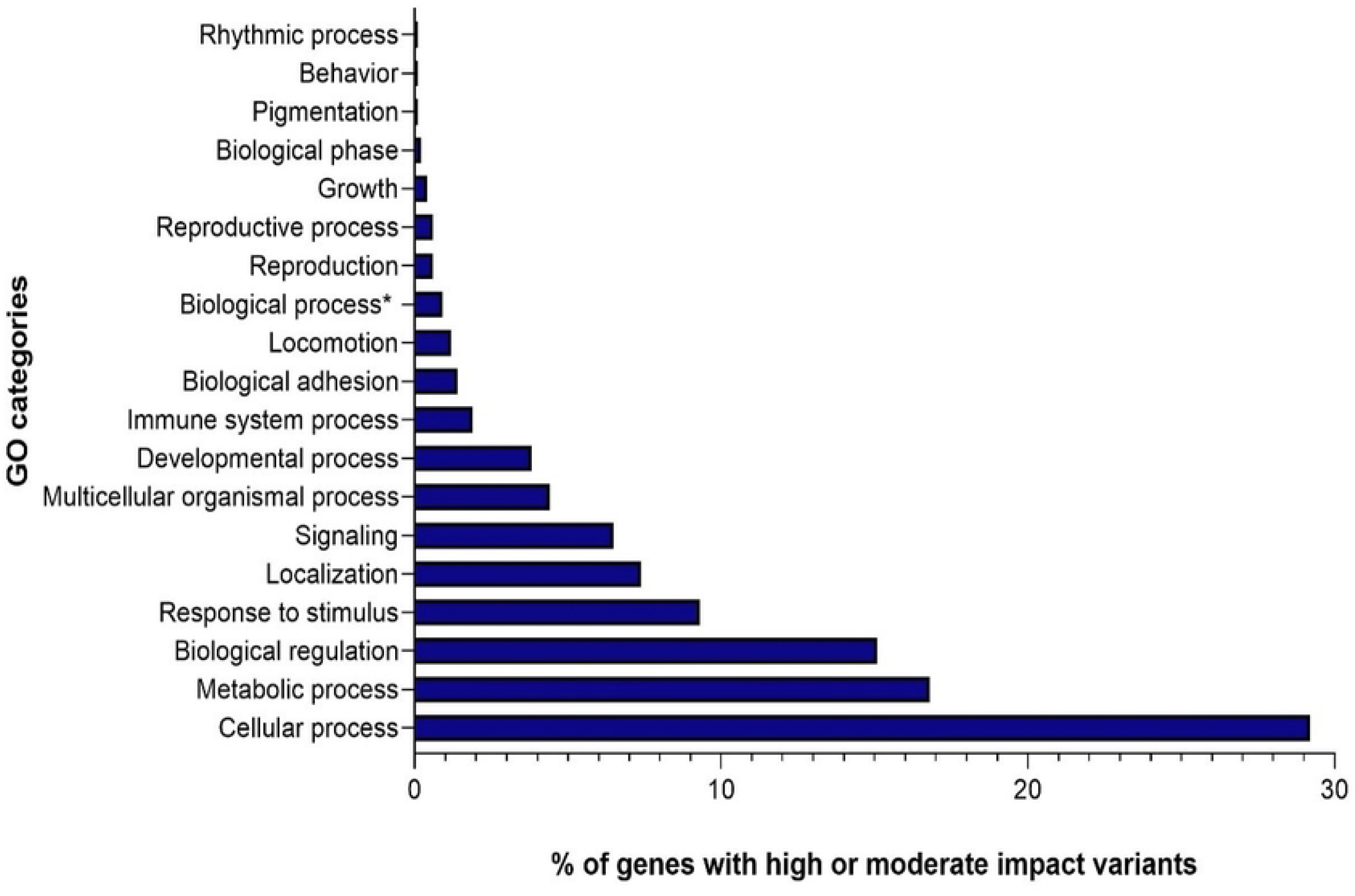
Distribution of functional categories for the percentage of genes containing high or moderate impact variants in Red Chittagong cattle (RCC).

The cellular process includes identified 75 significantly overrepresented GO terms in Red Chittagong cattle. Results GO analysis showed that a number of GO terms associated with immune functions were overrepresented such as Inflammatory response (GO:0006954), defense response (GO:0006952), leukocyte mediated immunity (GO:0002443), immune system process (GO:0002376), immune effector process (GO:0002252), regulation of immune response (GO:0050776), regulation of defense response (GO:0031347) and regulation of immune system process (GO:0002682). Immune functions related GO terms were also enriched in indicine cattle breeds from Pakistan and Finnish cattle breeds (36, 38). Overrepresented GO biological processes related to metabolic process were nitrogen compound metabolic process (GO:0006807), phosphate-containing compound metabolic process (GO:0006796), phosphorus metabolic process (GO:0006793), sulfur compound metabolic process (GO:0006790) and sulfur compound metabolic process (GO:0006790) (S1 Table).

A catalog of LoF variant with functional annotation would greatly aid gene prioritization by providing reference in disease studies (31, 39-41). In this study, eight KEGG terms were significantly enriched (FDR<0.10) for the genes containing LoF variants in RCC. Majority of them associated with immune response functional categories implicating the potential for RCC in investigating candidate variants for disease resistance or susceptibility.

### Variants identified in candidate gene associated with disease resistance

By manual curation of the annotated exonic variants, we identified 120 variants in 26 candidates for five diseases-foot and mouth disease (FMD), Mastitis, Parasite, para-tuberculosis and tick. Of the 120 variants, 50 were non-synonymous / frameshift (NS/FS) while 70 were synonymous/non-frameshift (SS/NFS) (S2 Table).

### Foot and mouth disease (FMD)

We observed nine polymorphisms in four genes associated with FMD. A frameshift deletion at 988 nucleotide position was observed in Bovine leukocyte antigen (*BOLA-A*) gene (Table 5). Eleven different alleles of *BOLA-DRB3* gene was reported in the Hariana cattle, of which three alleles were observed with protective immune response and three consistently ranked low for unprotected immune response for all the serotypes of FMDV (42). Similarly, Lei et al. (43) reported six RFLP patterns *of BoLA-DRB3* named AA, BB, CC, AB, AC and BC. CC and BC genotype were associated with resistance to FMD by contrast AA genotype was associated with susceptibility to FMD in Wanbei cattle.

**Table 4.** KEGG category enrichment (FDR <0.10) analysis for the genes containing loss-of-function (LoF) variants.

| Term | Fold Enrichment | p Value | FDR |
| --- | --- | --- | --- |
| AIG1-type G | 46.08 | 1.21E-07 | 4.08E-05 |
| AIG1 | 47.33 | 1.06E-07 | 4.52E-05 |
| MHC class I-like antigen recognition | 12.99 | 2.64E-06 | 5.64E-04 |
| Transferase | 2.41 | 1.37E-05 | 6.98E-04 |
| MHC classes I/II-like antigen recognition protein | 9.20 | 2.62E-05 | 0.003736 |
| Immunoglobulin-like fold | 2.38 | 3.85E-04 | 0.041122 |
| MHC class I-like antigen recognition-like | 13.11 | 5.52E-04 | 0.09261 |
| Basic and acidic residues | 1.46 | 8.24E-04 | 0.09261 |

**Table 5.** Functional variants identified in candidate genes for foot and mouth disease (FMD).

| Gene | Gene segment | Nucleotide change | Change in protein |
| --- | --- | --- | --- |
| *BOLA-A* | Exon5 | c.988delT | p.W330fs* |
| ***CCDC36*** | Exon8 | c.G1442A | p.R481Q |
| ***DDX10*** | Exon1 | c.G23T | p.R8L |
| ***DDX10*** | Exon1 | c.G84C | p.R28S |
| ***DDX10*** | Exon8 | c.C1087T | p.R363C |
| ***DDX10*** | Exon12 | c.A1420G | p.I474V |
| ***DDX10*** | Exon13 | c.C1862T | p.A621V |
| ***DDX10*** | Exon18 | c.C2510T | p.A837V |
| ***ITGB6*** | Exon14 | c.T2000C | p.F667S |

\* Frameshift deletion

In the present study, a nonsynonymous SNV was observed at the exon8 of the *CCDC36* gene (Table 5). Lee et al. (44) reported seven significantly FMD associated SNPs in spanning in the regions of *CCDC36* (4 in introns and 1 in an exon) and 2 were in intergenic regions between *CCDC36* and *C22H3orf62*. Their exonic SNP (rs110474439) in the *CCDC36* was a synonymous substitution, however, the nonsynonymous SNV observed in this study might of more interesting for functional study for FMD resistance. Lee et al. (44) also reported FMD associated SNP in an intron of the DEAD box polypeptide 10 (*DDX10*) gene, however, in this study we observed six nonsynonymous SNVs in 5 different exons of the *DDX10* gene. Based on the known function of the gene, we made inference that they might be involved in the functions related to innate immune responses.

A non-synonymous SNV was detected in the exon 14 of *ITGB6* gene. Singh et al. (45) in their study concluded that SNP G29A mutation in the 5 UTR of the *ITGB6* gene (chromosome 2) associated with resistance to FMD infection in the zebu cattle.

### Mastitis

Seven non-synonymous SNVs were detected in four mastitis associated genes (Table 6). Vitenberga-Verza et al. (46) showed that immunoreactivity was more pronounced for IL-4 in mastitis cases. We identified a non-synonymous mutation in the *IL4* gene that coding for IL4. We detected four missense mutations in three different exons of BRCA1 gene. Using combined genotype analysis of three SNPs -G22231T, T25025A and C28300A in bovine *BRCA1* gene Yuan et al. (47) showed association of BBDDFF genotype with the highest somatic cell score (SCS) that indicated mastitis susceptibility. However, AACCEE genotype associated with the lowest SCS was favorable for the mastitis resistance. Magotra et al. (48) also reported association of *BRCA1* gene SNPs with mastitis susceptibility and resistance.

**Table 6.**
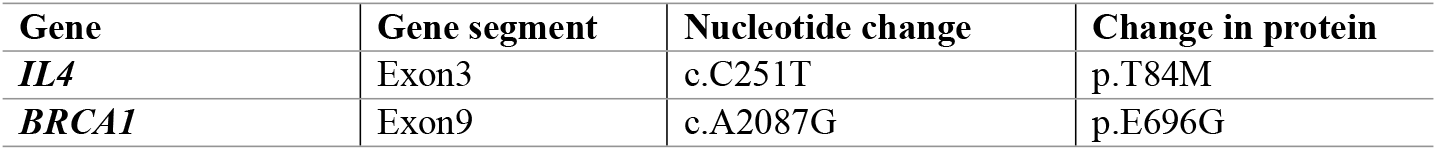
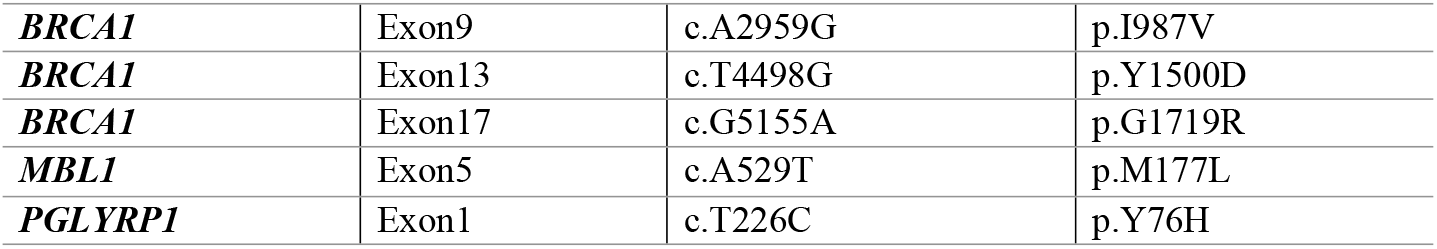
Functional variants identified in candidate genes for mastitis.

We observed a non-synonymous SNV was observed at exon5 of the *MBL1* gene. A reported significant association between a SNP (g.2651G>A) in *MBL1* gene and SCS, suggesting a possible role of this SNP in the host response against mastitis (49). Zabolewicz et al. (50) showed relationship between polymorphism within Peptidoglycan Recognition Protein 1 gene (*PGLYRP1*) and somatic cell counts in milk of Holstein cows. In this study we also observed a SNV in the exon 1 of the *PGLYRP1* gene.

### Candidate variants for parasite resistance

Twelve nonsynonymous SNVs were identified in four genes associated with parasite resistance (Table 7). A genome-wide associations and functional gene analyses for endoparasite resistance in an endangered population of native German Black Pied cattle revealed *ALCAM* gene is a potential candidate gen for endoparasite resistance trait (28). In this we detected a missense mutation at 58 nucleotide position of the *ALCAM* gene (Table 10).

**Table 7.** Nonsynonymous SNVs identified in candidate genes for parasite resistance.

| Gene | Gene segment | Nucleotide change | Change in protein |
| --- | --- | --- | --- |
| *ALCAM* | exon1 | c.A58G | p.T20A |
| *ARHGAP15* | exon10 | c.G873T | p.K291N |
| *CATHL1* | exon4 | c.A451T | p.I151L |
| *CATHL1* | exon4 | c.A383G | p.Q128R |
| *CATHL1* | exon4 | c.A451T | p.I151L |
| *CATHL3* | exon4 | c.T474G | p.I158M |
| *CATHL3* | exon4 | c.T473G | p.I158S |
| *CDH2* | exon2 | c.G72T | p.E24D |
| *CDH2* | exon10 | c.G1543A | p.V515M |
| *CDH2* | exon11 | c.G1702A | p.A568T |
| *CDH2* | exon11 | c.C1703A | p.A568D |
| *CDH2* | exon11 | c.A1708G | p.I570V |

A genetic and expression analysis of African cattle identified *ARHGAP15* as a candidate gene in pathways responding to *Trypanosoma congolense* infection Noyesa et al. (51). Present study detected a G to T substitution resulting a Lysine to asparagine change in ARHGAP15 protein. *CATHL1* gene was observed to carry five nonsynonymous SNVs in the sequenced bull. Whole genome sequencing of Gir cattle revealed a selective sweep that harbours several genes belonging to the cathelicidin gene family, such as *CAMP, CATHL1, CATHL2*, and *CATHL3*, which are related to pathogen- and parasite-resistance (52). Five variants were found in three exons of the *CDH2* gene. May et al., (28) showed that *CDH2* gene had a significance association with endoparasite resistance trait.

## Conclusion

The present study is the first to perform whole genome sequencing and detect genetic variations in RCC. Sequencing with a mean depth of 31X coverage leads to identifying 3324621 novel variants. Functional annotation of detected variants revealed 24575 homozygous exonic variants, including non-synonymous SNVs, stop gain frameshift INDELs that will be a resource for establishing breed-specific genotyping arrays. In addition, genetic differences between RCC and taurine reference sequences identified in the present study provide valuable markers for future association studies for economic phenotypes. Potentially functional variants were identified in genes associated with disease resistance, such as *BoLA-DRB3, BRCA1, CCDC36, CDH2, DDX10, IL4, ITGB6, MBL1, and PGLYRP1*. Moreover, variants were detected in genes that reported parasite resistance, such as *ALCAM, ARHGAP15, CATHL1*, and *CDH2*. This catalog of detected variants will be helpful in future studies to understand genetic variation underlying disease resistance in indicine breeds.

## Acknowledgments

The authors acknowledge the computing resources of the research school of Biology, Australian National University and thank Associate Prof. Alexander Mikheyev for his support for computational facilities. The authors also acknowledge BRAC Artificial Insemination Enterprise for providing the semen sample for this study. Finally, the authors thank the Poultry Research and Training Center, Chattogram, for supporting wet lab facilities.

## References

1. Le Roex N, Noyes H, Brass A, Bradley DG, Kemp SJ, Kay S, et al. Novel SNP discovery in African buffalo, Syncerus caffer, using high-throughput sequencing. PloS one. 2012;7(11):e48792.

2. Mullen MP, Creevey CJ, Berry DP, McCabe MS, Magee DA, Howard DJ, et al. Polymorphism discovery and allele frequency estimation using high-throughput DNA sequencing of target-enriched pooled DNA samples. BMC genomics. 2012;13(1):1–12.

3. Canavez FC, Luche DD, Stothard P, Leite KR, Sousa-Canavez JM, Plastow G, et al. Genome sequence and assembly of Bos indicus. Journal of Heredity. 2012;103(3):342–8.

4. Rosen BD, Bickhart DM, Schnabel RD, Koren S, Elsik CG, Tseng E, et al. De novo assembly of the cattle reference genome with single-molecule sequencing. Gigascience. 2020;9(3):giaa021.

5. Chen N, Fu W, Zhao J, Shen J, Chen Q, Zheng Z, et al. BGVD: an integrated database for bovine sequencing variations and selective signatures. Genomics, Proteomics & Bioinformatics. 2020;18(2):186–93.

6. Das S, Bhuiyan A, Begum N, Habib M, Arefin T. Fertility and parasitic infestation of Red Chittagong cattle. Bangladesh Veterinarian. 2010;27(2):74–81.

7. Habib M, Bhuiyan A, Amin M. Reproductive performance of Red Chittagong cattle in a nucleus herd. Bangladesh Journal of Animal Science. 2010;39(1-2):9–19.

8. Halim M, Kashem M, Ahmed J, Hossain M. Economic analysis of Red Chittagong Cattle farming system in some selected areas of Chittagong district. Journal of the Bangladesh Agricultural University. 2010;8(2):271–6.

9. Khan M, Miah G, Huque K, Khatun M, Das A. Economic and genetic evaluations of different dairy cattle breeds under rural conditions in Bangladesh. Livest Res Rural Devel. 2011;24(01).

10. Khan M, Miah G, Khatun M, Das A. Economic values for different economic traits of Red Chittagong cow. Indian Journal of Animal Sciences. 2010;80(11):1138.

11. Ahmed R, Biswas PK, Barua M, Alim MA, Islam K, Islam MZ. Prevalence of gastrointestinal parasitism of cattle in Banskhali upazilla, Chittagong, Bangladesh. Journal of Advanced Veterinary and Animal Research. 2015;2(4):484–8.

12. Badruzzaman A, Siddiqui MSI, Faruk MO, Lucky NS, Zinnah MA, Hossain FMA, et al. Prevalence of infectious and non-infectious diseases in cattle population in Chittagong district of Bangladesh. International Journal of Biological Research. 2015;3(1):1–4.

13. Siddiki A, Uddin M, Hasan M, Hossain M, Rahman M, Das B, et al. Coproscopic and Haematological Approaches to Determine the Prevalence of Helminthiasis and Protozoan Diseases of Red Chittagong Cattle (RCC) Breed in Bangladesh. Pakistan Veterinary Journal. 2010;30(1).

14. Ferdous F, Choudhury M, Faruque M, Hossain M, Bhuiyan A. Genetic evaluation of Red Chittagong cattle in Bangladesh. SAARC Journal of Agriculture. 2019;17(2):141–54.

15. Nahar S, Islam Afmf, Hoque MA, Bhuiyan Akfh. Animal performance of indigenous Red Chittagong cattle in Bangladesh. Acta Scientiarum Animal Sciences. 2016;38:177–82.

16. Bhuiyan M, Bhuiyan A, Yoon D, Jeon J, Park C, Lee J. Mitochondrial DNA diversity and origin of Red Chittagong cattle. Asian-Australasian Journal of Animal Sciences. 2007;20(10):1478–84.

17. Edea Z, Bhuiyan M, Dessie T, Rothschild M, Dadi H, Kim K. Genome-wide genetic diversity, population structure and admixture analysis in African and Asian cattle breeds. Animal. 2015;9(2):218–26.

18. Mandefro A, Sisay T, Edea Z, Uzzaman MR, Kim K-S, Dadi H. Genetic assessment of BoLA-DRB3 polymorphisms by comparing Bangladesh, Ethiopian, and Korean cattle. Journal of Animal Science and Technology. 2021;63(2):248.

19. Mufti MMR, Mostari MP, Deb GK, Nahar K, Huque KS. Genetic diversity of Red Chittagong Cattle using randomly amplified polymorphic DNA markers. American Journal of Animal and Veterinary Sciences. 2009;4(1):1–5.

20. Uzzaman MR, Edea Z, Bhuiyan MSA, Walker J, Bhuiyan A, Kim K-S. Genome-wide single nucleotide polymorphism analyses reveal genetic diversity and structure of wild and domestic cattle in Bangladesh. Asian-Australasian Journal of Animal Sciences. 2014;27(10):1381.

21. Chen Y, Chen Y, Shi C, Huang Z, Zhang Y, Li S, et al. SOAPnuke: a MapReduce acceleration-supported software for integrated quality control and preprocessing of high-throughput sequencing data. Gigascience. 2018;7(1):gix120.

22. Andrews S. FastQC: a quality control tool for high throughput sequence data. Babraham Bioinformatics, Babraham Institute, Cambridge, United Kingdom; 2010.

23. Li H, Durbin R. Fast and accurate long-read alignment with Burrows–Wheeler transform. Bioinformatics. 2010;26(5):589–95.

24. Li H, Handsaker B, Wysoker A, Fennell T, Ruan J, Homer N, et al. The sequence alignment/map format and SAMtools. Bioinformatics. 2009;25(16):2078–9.

25. Li H. A statistical framework for SNP calling, mutation discovery, association mapping and population genetical parameter estimation from sequencing data. Bioinformatics. 2011;27(21):2987–93.

26. Wang K, Li M, Hakonarson H. ANNOVAR: functional annotation of genetic variants from high-throughput sequencing data. Nucleic acids research. 2010;38(16):e164–e.

27. Mi H, Muruganujan A, Huang X, Ebert D, Mills C, Guo X, et al. Protocol Update for large-scale genome and gene function analysis with the PANTHER classification system (v. 14.0). Nature protocols. 2019;14(3):703–21.

28. May K, Scheper C, Brügemann K, Yin T, Strube C, Korkuć P, et al. Genome-wide associations and functional gene analyses for endoparasite resistance in an endangered population of native German Black Pied cattle. BMC genomics. 2019;20(1):1–15.

29. Prajapati B, Gupta J, Pandey D, Parmar G, Chaudhari J. Molecular markers for resistance against infectious diseases of economic importance. Veterinary world. 2017;10(1):112.

30. Singh U, Deb R, Alyethodi RR, Alex R, Kumar S, Chakraborty S, et al. Molecular markers and their applications in cattle genetic research: A review. Biomarkers and Genomic medicine. 2014;6(2):49–58.

31. Das A, Panitz F, Gregersen VR, Bendixen C, Holm L-E. Deep sequencing of Danish Holstein dairy cattle for variant detection and insight into potential loss-of-function variants in protein coding genes. BMC genomics. 2015;16(1):1–12.

32. Eck SH, Benet-Pagès A, Flisikowski K, Meitinger T, Fries R, Strom TM. Whole genome sequencing of a single Bos taurusanimal for single nucleotide polymorphism discovery. Genome biology. 2009;10(8):1–8.

33. Stothard P, Choi J-W, Basu U, Sumner-Thomson JM, Meng Y, Liao X, et al. Whole genome resequencing of black Angus and Holstein cattle for SNP and CNV discovery. BMC genomics. 2011;12(1):1–14.

34. Zhan B, Fadista J, Thomsen B, Hedegaard J, Panitz F, Bendixen C. Global assessment of genomic variation in cattle by genome resequencing and high-throughput genotyping. BMC genomics. 2011;12(1):1–20.

35. Zwane AA, Schnabel RD, Hoff J, Choudhury A, Makgahlela ML, Maiwashe A, et al. Genome-wide SNP discovery in indigenous cattle breeds of South Africa. Frontiers in Genetics. 2019;10:273.

36. Iqbal N, Liu X, Yang T, Huang Z, Hanif Q, Asif M, et al. Genomic variants identified from whole-genome resequencing of indicine cattle breeds from Pakistan. PLoS One. 2019;14(4):e0215065.

37. Rosse IC, Assis JG, Oliveira FS, Leite LR, Araujo F, Zerlotini A, et al. Whole genome sequencing of Guzerá cattle reveals genetic variants in candidate genes for production, disease resistance, and heat tolerance. Mammalian genome. 2017;28(1):66–80.

38. Weldenegodguad M, Popov R, Pokharel K, Ammosov I, Ming Y, Ivanova Z, et al. Whole-genome sequencing of three native cattle breeds originating from the northernmost cattle farming regions. Frontiers in genetics. 2019;9:728.

39. MacArthur DG, Tyler-Smith C. Loss-of-function variants in the genomes of healthy humans. Human molecular genetics. 2010;19(R2):R125–R30.

40. TG, HDL Working Group of the Exome Sequencing Project NH, Lung,, Institute B. Loss-of-function mutations in APOC3, triglycerides, and coronary disease. New England Journal of Medicine. 2014;371(1):22–31.

41. Xu D, Gokcumen O, Khurana E. Loss-of-function tolerance of enhancers in the human genome. PLoS Genetics. 2020;16(4):e1008663.

42. Gowane G, Sharma A, Sankar M, Narayanan K, Das B, Subramaniam S, et al. Association of BoLA DRB3 alleles with variability in immune response among the crossbred cattle vaccinated for foot-and-mouth disease (FMD). Research in veterinary science. 2013;95(1):156–63.

43. Lei W, Liang Q, Jing L, Wang C, Wu X, He H. BoLA-DRB3 gene polymorphism and FMD resistance or susceptibility in Wanbei cattle. Molecular Biology Reports. 2012;39(9):9203–9.

44. Lee B-Y, Lee K-N, Lee T, Park J-H, Kim S-M, Lee H-S, et al. Bovine genome-wide association study for genetic elements to resist the infection of foot-and-mouth disease in the field. Asian-Australasian journal of animal sciences. 2015;28(2):166.

45. Singh R, Deb R, Singh U, Alex R, Kumar S, Chakraborti S, et al. Development of a tetra-primer ARMS PCR-based assay for detection of a novel single-nucleotide polymorphism in the 5′ untranslated region of the bovine ITGB6 receptor gene associated with foot-and-mouth disease susceptibility in cattle. Archives of virology. 2014;159(12):3385–9.

46. Vitenberga-Verza Z, Pilmane M, Šerstņova K, Melderis I, Gontar Ł, Kochański M, et al. Identification of Inflammatory and Regulatory Cytokines IL-1α-, IL-4-, IL-6-, IL-12-, IL-13-, IL-17A-, TNF-α-, and IFN-γ-Producing Cells in the Milk of Dairy Cows with Subclinical and Clinical Mastitis. Pathogens. 2022;11(3):372.

47. Yuan Z, Chu G, Dan Y, Li J, Zhang L, Gao X, et al. BRCA1: a new candidate gene for bovine mastitis and its association analysis between single nucleotide polymorphisms and milk somatic cell score. Molecular biology reports. 2012;39(6):6625–31.

48. Magotra A, Gupta I, Ahmad T, Alex R. Polymorphism in DNA repair gene BRCA1 associated with clinical mastitis and production traits in indigenous dairy cattle. Research in Veterinary Science. 2020;133:194–201.

49. Wang C, Liu M, Li Q, Ju Z, Huang J, Li J, et al. Three novel single-nucleotide polymorphisms of MBL1 gene in Chinese native cattle and their associations with milk performance traits. Veterinary immunology and immunopathology. 2011;139(2-4):229–36.

50. Zabolewicz T, Puckowska P, Brym P, Oleński K, Kamiński S. Relationship between polymorphism within Peptidoglycan Recognition Protein 1 gene (PGLYRP1) and somatic cell counts in milk of Holstein cows. Annals of Animal Science. 2022;22(2):593–9.

51. Noyes H, Brass A, Obara I, Anderson S, Archibald AL, Bradley DG, et al. Genetic and expression analysis of cattle identifies candidate genes in pathways responding to Trypanosoma congolense infection. Proceedings of the National Academy of Sciences. 2011;108(22):9304–9.

52. Liao X, Peng F, Forni S, McLaren D, Plastow G, Stothard P. Whole genome sequencing of Gir cattle for identifying polymorphisms and loci under selection. Genome. 2013;56(10):592–8.

